# A single mutation in dairy cow-associated H5N1 viruses increases receptor binding breadth

**DOI:** 10.1101/2024.06.22.600211

**Authors:** Marina R. Good, Wei Ji, Monica L. Fernández-Quintero, Andrew B. Ward, Jenna J. Guthmiller

**Author notes:** Contributed equally (Marina R. Good, Wei Ji).

## Abstract

Clade 2.3.4.4b H5N1 is causing an unprecedented outbreak in dairy cows in the United States. To understand if recent H5N1 viruses are changing their receptor use, we screened recombinant hemagglutinin (HA) from historical and recent 2.3.4.4b H5N1 viruses for binding to distinct glycans bearing terminal sialic acids. We found that H5 from A/Texas/37/2024, an isolate from the dairy cow outbreak, has increased binding breadth to glycans bearing terminal α2,3 sialic acids, the avian receptor, compared to historical and recent 2.3.4.4b H5N1 viruses. We did not observe any binding to α2,6 sialic acids, the receptor used by human seasonal influenza viruses. We identified a single mutation outside of the receptor binding site, T199I, was responsible for increased binding breadth, as it increased receptor binding site flexibility. Together, these data show recent H5N1 viruses are evolving increased receptor binding breadth which could impact the host range and cell types infected with H5N1.

## INTRODUCTION

Since 2021, clade 2.3.4.4b H5N1 viruses, a highly pathogenic avian influenza virus, have been causing a worldwide outbreak in wild bird populations, with reported cases on six continents. Numerous H5N1 spillover events in domestic animals, including poultry and minks, have led to massive culling events^1,2^. Moreover, H5N1 spillover into wild mammals, including aquatic and scavenger mammals, have been reported since 2022^3^. In March 2024, the United States Department of Agriculture reported an outbreak of H5N1 in domestic dairy cattle^4^. Since then, H5N1 has expanded to 12 states with over 100 farms affected^5^. H5N1 viruses from dairy cows have spilled over into domestic felines, alpacas, poultry, and house mice^6,7^. Importantly, H5N1 viruses from the ongoing outbreak in dairy cattle have led to three confirmed human infections, with two cases causing conjunctivitis and the third case cause mild respiratory symptoms^8,9^.

H5N1 infection in dairy cows is largely restricted to the mammary tissue, leading to clinical manifestations of mastitis including reductions in milk production, milk discoloration, and increased milk thickness^10^. Analysis of infectious virus revealed titers ranging from 10^4^-10^9^ tissue culture infectious dose 50 (TCID_50_)^11,12^. One in five retail milk samples within the United States (US) has detectable virus by PCR, although viable virus has not been recovered from these samples^13^. Moreover, pasteurization is an effective method to kill H5N1 viruses^11,14^. It remains unclear how H5N1 is being transmitted between cows and different hosts, although transmission is linked to raw milk consumption or exposure.

Avian influenza viruses, including H5N1, preferentially bind glycans bearing terminal α2,3 sialic acids^15^. In contrast, influenza viruses that cause seasonal influenza outbreaks in humans prefer glycans bearing terminal α2,6 sialic acids^16^. The influenza virus preference for α2,3 or α2,6 sialic acid linkages creates a major species barrier for avian influenza viruses to spill over into humans. Two recent studies show that dairy cow mammary tissue, and particularly the mammary alveoli, has abundant α2,3 sialic acid linked glycans^17,18^. Moreover, dairy cow mammary tissues also have α2,6 sialic acid linked glycans^17,18^, suggesting dairy cow mammary glands could be a site of viral evolution to adapt H5N1 to human-like receptors.

In this study, we investigated if recent H5N1 viruses are evolving their receptor binding specificities. We identified that H5 from the ongoing dairy cow outbreak has increased binding breadth to backbone glycans bearing α2,3 sialic acids relative to other H5N1 viruses, which was linked to a single mutation near, but not within, the receptor binding site (RBS). I199 emerged in late 2023, before the onset of the ongoing dairy cow outbreak, and is now the dominant amino acid at this residue in North American isolates. Our study indicates a single mutation near the RBS expands the types of backbone gylcans bound by H5, which could imply an increase in cell, tissue and host tropisms.

## RESULTS

### HA from circulating H5N1 in dairy cows is phylogenetically distinct

The phylogenetic analysis of HA genetic sequences from 94 H5Nx viruses representing various clades and ancestral and human H5N1 sequences revealed distinct branching patterns (**Fig. 1**; **Extended Data Table 1**). Human H5N1 sequences from Vietnam (2004) and Indonesia (2005) were closely related but formed separate branches from the 2.3.4.4 clusters, bridging the evolutionary paths between the ancestral A/Goose/Guangdong/1996 sequence and the 2.3.4.4 clades (**Fig. 1**). The 2.3.4.4 clade is further subdivided into distinct subclusters (2.3.4.4b, 2.3.4.4c, 2.3.4.4e, 2.3.4.4g, 2.3.4.4h), highlighting the diversity and evolutionary progression of H5Nx across various regions. The 2.3.4.4b clade represents the dominant H5N1 viruses globally since 2021^19–21^. 2.3.4.4.b H5N1 viruses segregated based on continent(s), with virus isolates from Eurasia and Africa more closely related to each other than to viruses from North and South America, which was independent of isolation date (**Fig. 1**). Moreover, viruses from North and South America are more closely related to each other than they are to viruses from Eurasia and Africa (**Fig. 1**). Within the Americas branches, the cattle-derived H5N1 viruses form a distinct group within the 2.3.4.4b clade (**Fig. 1**). The new group shows a cluster of H5N1 strains isolated from various hosts in the United States in 2024, including dairy cows, domestic cats, raccoons, skunks, mountain lions, and several bird species. The clustering of sequences from the recent dairy cow outbreak indicates a common ancestor and suggests a potential transmission link between these species. Together, these data demonstrate that the recent outbreak of H5N1 in dairy cows is distinct from other circulating 2.3.4.4b H5N1 viruses.

**Fig. 1:**
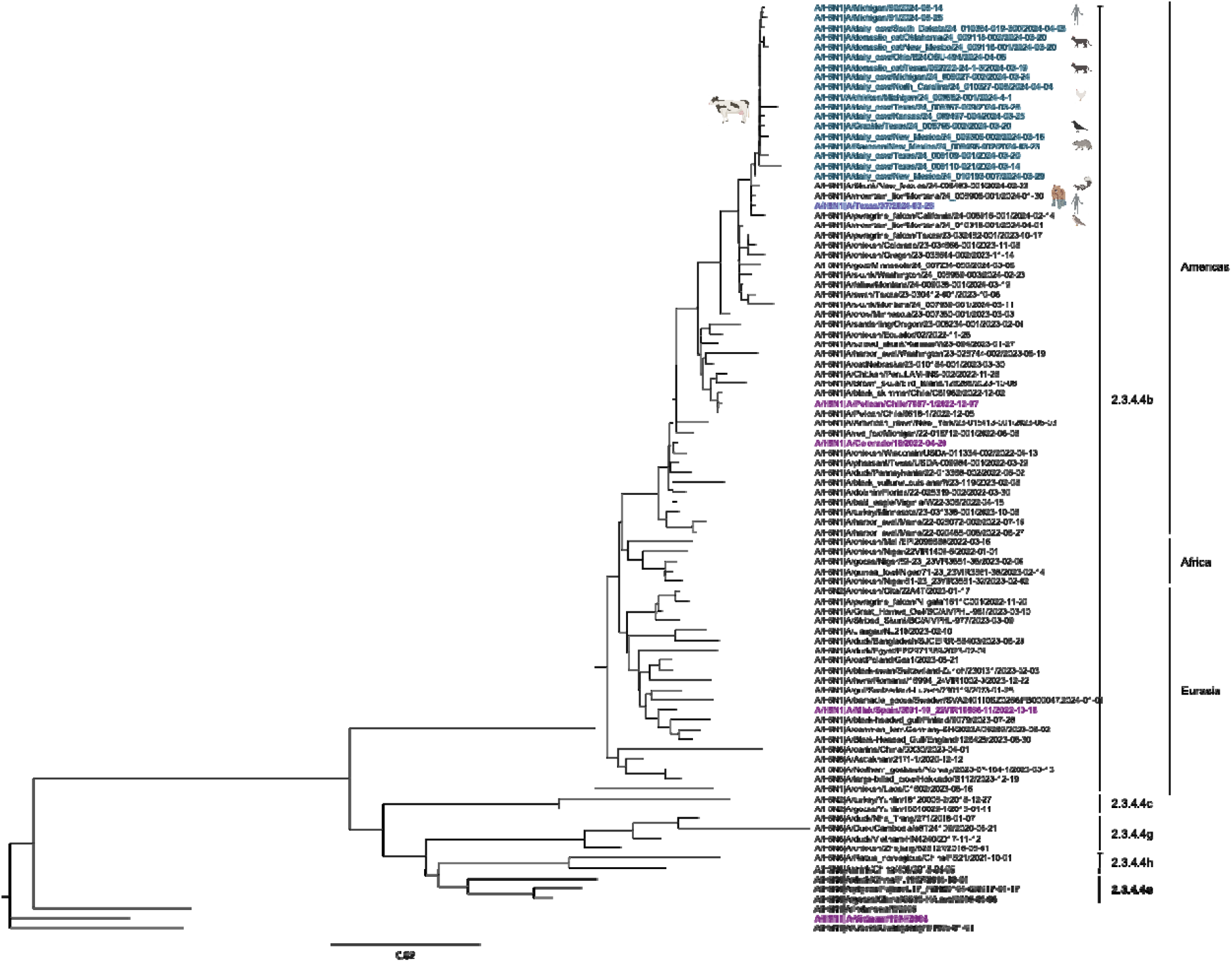
Phylogenetic tree of highly pathogenic avian H5N1. The Neighbor-Joining (NJ) phylogenetic analysis of 94 hemagglutinin gene sequences. Clade 2.3.4.4 is subdivided into distinct subclades, including 2.3.4.4e, 2.3.4.4h, 2.3.4.4g, 2.3.4.4c, and the currently dominant 2.3.4.4b clade. The new group belonging to clade 2.3.4.4b includes strains isolated from domestic dairy cows and humans and animals linked to H5N1-positive dairy farms (highlighted with a blue region and animal symbols) in the United States in 2024. Distinct clades of the virus are labeled on the right side of the figure. Tips are labeled with H5Nx strain names, host species, and isolation dates. Nodes represent inferred common ancestors of the grouped tips. Branch lengths are proportional to the number of nucleotide substitutions per site, indicating the divergence between nodes.

### Dairy cow-related H5 has increased glycan binding breadth

To understand if recent H5 has changed its receptor binding specificity, we tested recombinant H5 (rH5) from an ancestral H5N1 virus, 2.3.4.4b H5N1 viruses from 2022, and a recent H5 isolated from dairy farm worker (A/Texas/37/2024) on a N-acetylneuraminic (Neu5Ac) and N-glyconeuraminic acid (Neu5Gc) glycan microarray (**Extended Data Table 2**). This microarray includes an array of distinct glycans with terminal sialic acids of both the α2,3 and α2,6 Neu5Ac linkages, which correspond to the receptors for avian and human influenza viruses, respectively. This microarray includes glycans with distinct branches, with most glycans incorporating a single branch with α2,3 or α2,6 Neu5Ac linkage and a second branch of varying lengths and compositions (**Extended Data Table 2**). We observed ancestral rH5 from A/Vietnam/1204/2004 exhibited a dominant preference for α2,3 linked lactosamine glycans (**Fig. 2A**). In contrast, rH5 from A/Colorado/18/2022, which was isolated from a human involved in culling 2.3.4.4b H5N1 infected poultry, exhibited restricted binding to 3’ sialyl Lewis X glycans. Other isolates from 2022 2.3.4.4b H5N1 viruses revealed expanded binding breadth to 3’ sialyl Lewis X and α2,3 linked lactosamine glycans (**Fig. 2A**). Compared to other 2.3.4.4b rH5s, A/Texas/37/2024 has gained further binding breadth to nearly all α2,3 sialic linked lactosamine glycans, including those with asymmetrical branches (**Fig. 2A**). Moreover, A/Texas/37/2024 had augmented binding signal for 3’ sialyl Lewis X glycans relative other 2.3.4.4b H5 viruses and to α2,3 sialic acid-linked lactosamine glycans relative to A/Vietnam/1204/2004 (**Fig. 2A; Extended Data Fig. 1A-E**). Importantly, we did not observe any binding to glycans bearing only terminal α2,6 sialic acids, indicating recent H5N1 viruses have not gained binding affinity to receptors used by human seasonal influenza virus subtypes (**Extended Data Fig. 1A-E**).

**Fig. 2:**
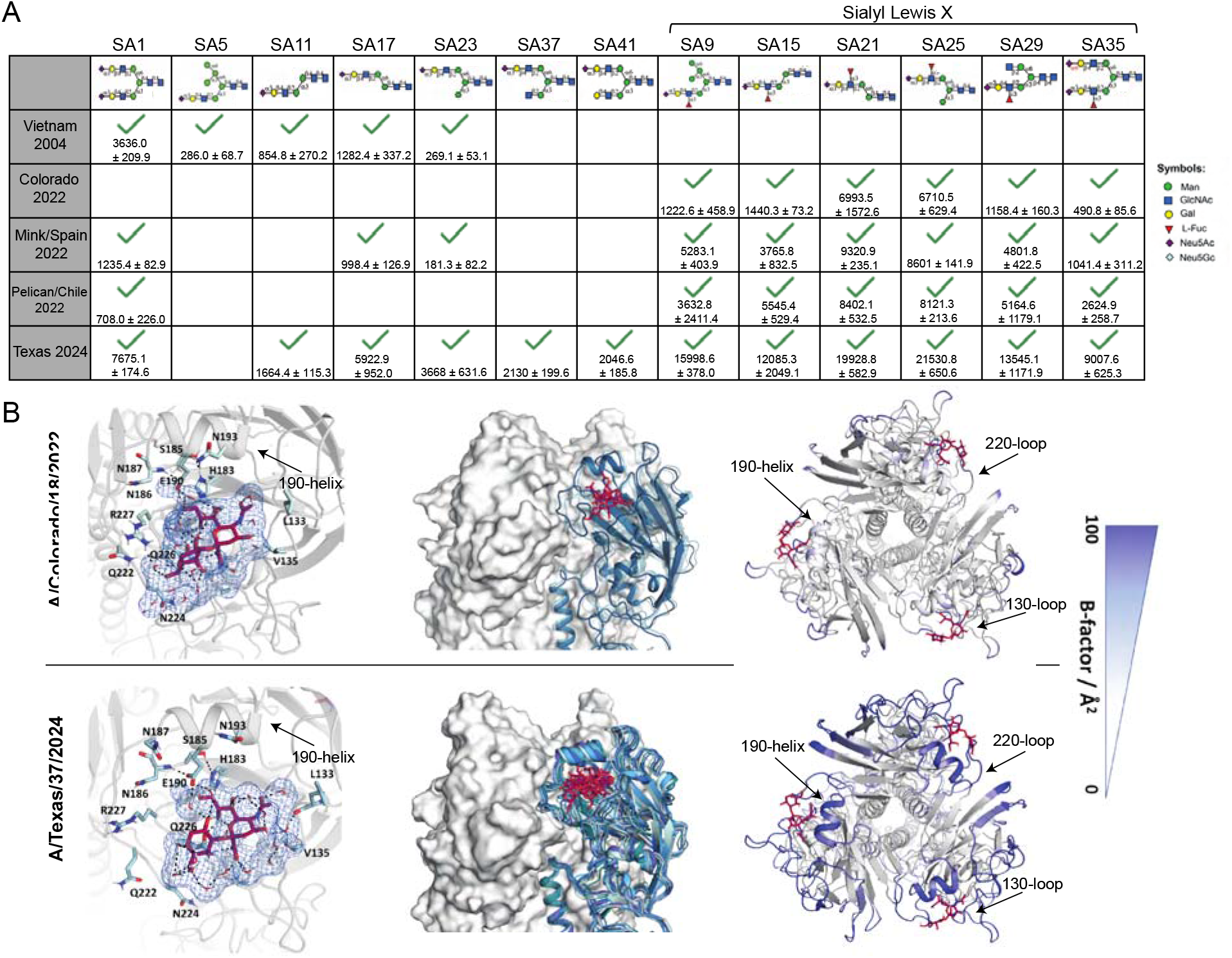
A dairy cow associated H5N1 virus exhibits increased glycan binding breadth. (A) rH5 binding to distinct Neu5Ac glycans. Green checkmarks indicate a positive binding result for the corresponding glycan. Normalized relative fluorescence unit (RFU) values ± standard deviation are indicated below checkmarks. Value above each glycan indicates the glycan number in the array. (B-D) MD simulations of A/Colorado/18/2022 and A/Texas/37/2024 to characterize the LSTa binding site properties. (B) Representative structure obtained from MD simulations showing interactions of LSTa (burgundy) with A/Colorado/18/2022 and A/Texas/37/2024 in solution. (C) Conformational states of A/Colorado/18/2022 and A/Texas/37/2024 binding to LSTa. Each shade of blue represents a distinct confirmation. (D) Residue-wise B-factor, as a measure of flexibility, mapped on the respective A/Colorado/18/2022 and A/Texas/37/2024 structure.

To understand the mechanism of how A/Texas/37/2024 rH5 has gained increased binding breadth, we performed molecular dynamics (MD) simulations. Sequences of rH5 from A/Texas/37/2024 and A/Colorado/18/2022 were used to model binding to LSTa, an α2,3 sialic acid avian analog receptor (**Fig. 2B**). We identified similar residues of both A/Colorado/18/2022 and A/Texas/37/2024 involved in LSTa binding (**Fig. 2B**). Additionally, we observed a strong water-mediated, hydrogen bond network of LSTa with both A/Texas/37/2024 and A/Colorado/18/2022. In particular, we found a long-lasting water-mediated hydrogen bond of LSTa with residue E190 (**Fig. 2B**), an important residue for mediating α2,3 sialic acid receptor specificity^22^. A/Texas/37/2024 exhibited more variability in the receptor binding site (RBS), adopting 9 different conformations (**Fig. 2C**). In contrast A/Colorado/18/2022 reveals a more stable RBS with only 3 conformational states (**Fig. 2C**). To quantify the differences in flexibility, we mapped the B-factor obtained from the MD simulations onto the structure of A/Colorado/18/2022 and A/Texas/372/2024 in complex with LSTa. A/Texas/37/2024 exhibited overall a dramatically higher B-factor. This is particularly pronounced in the 190-helix, whereas A/Colorado/18/2022 revealed an overall rather low B-factor (**Fig. 2D**). Thus, our findings suggest that the expanded binding breadth to glycans even bearing terminal α2,3 sialic acids, is mediated by an increased flexibility within the RBS of A/Texas/37/2024.

### H5N1 viruses circulating in the Americas have acquired mutations near the RBS

Several mutations have arisen in 2.3.4.4b viruses since 2022, particularly at L111M, T199I, and V214A (**Fig. 3A**). Notably, these three mutations lie outside of the traditional RBS, which is comprised of the 130-loop, 190-helix, and 220-loop^23^. Analysis of these mutations based on location and outbreak revealed that all three mutations are specific to H5N1 viruses in the Americas (**Fig. 3B**). Importantly, we observed T199I was found in all H5N1 viruses from the dairy cow outbreak, as well as some H5N1 viruses circulating in the Americas not related to the ongoing dairy cow outbreak (**Fig. 3B**). T199I is the only amino acid difference between A/Texas/37/2024 and A/pelican/Chile/7087-1/2022 (**Fig. 3C-D**). Importantly, HAs from the ongoing dairy cow outbreak are highly conserved, as the amino acid sequence of A/Texas/37/2024 and A/Michigan/90/2015 are identical (**Fig. 3C**). Structurally, position 199 is located on the backside of the 190-helix (**Fig. 3A**). We observed that the T199I mutation arose in the second half of 2023, with I199 becoming dominate by November 2023 (**Fig. 3D-E**). Notably, A/Texas/37/2024 HA NT sequence clusters more closely with an HA sequence collected from a mountain lion, also known as a cougar (*Puma concolor*) in Montana than sequences related to the ongoing dairy outbreak (**Fig. 3E**). Interestingly, the mountain lion isolate was collected in January 2024 and it was recently proposed that the dairy cow outbreak has been ongoing since late 2023^24^. These data would suggest a closely related common ancestor from the mountain lion case to the ongoing dairy cow outbreak. Together, these data show that H5N1 viruses in the Americas have accumulated mutations within the RBS, with T199I being the only mutation specific to the ongoing dairy cow outbreak.

**Fig. 3:**
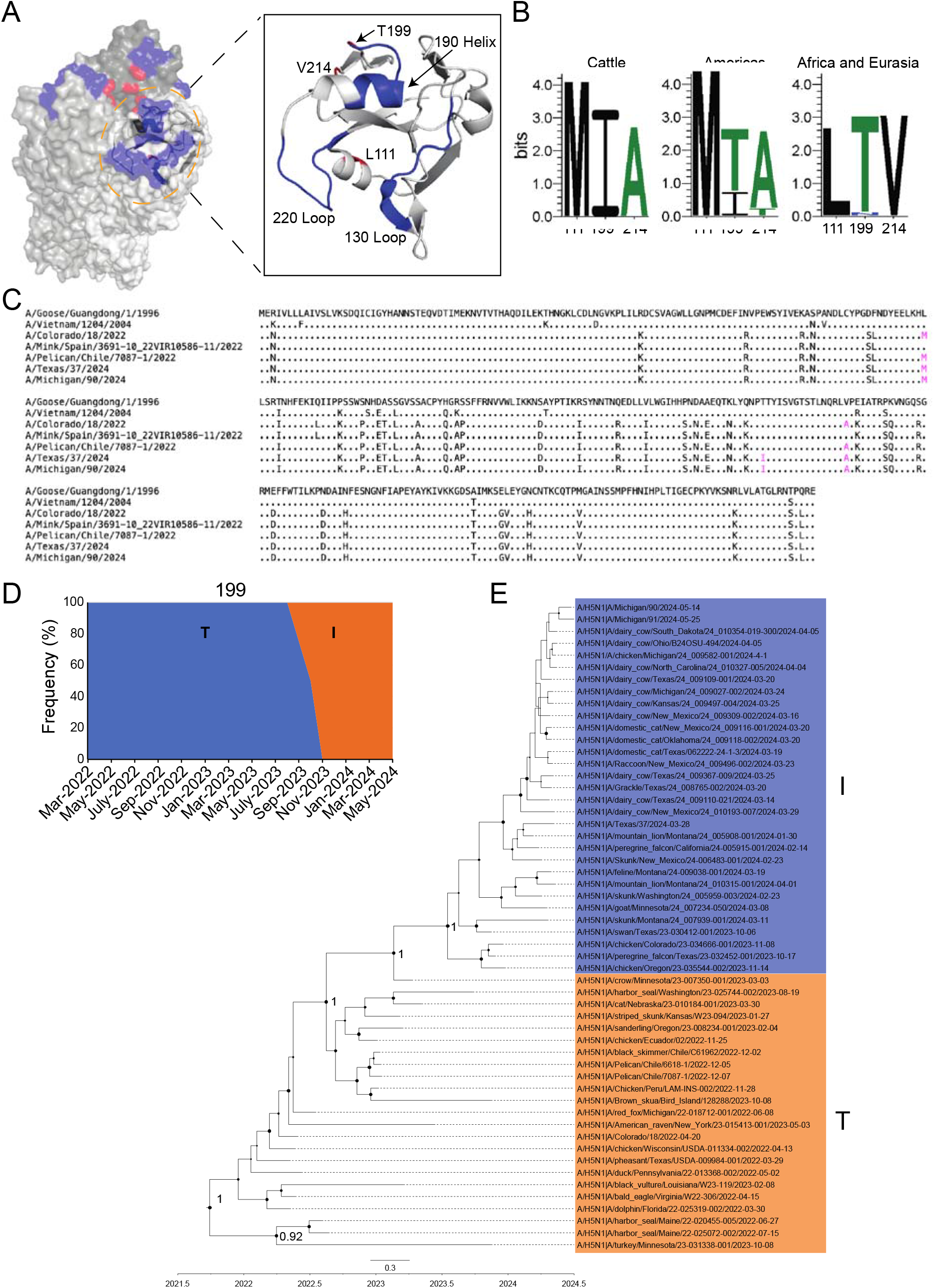
2.3.4.4b H5N1 viruses in the Americas recently acquired T199I. (A) Structural depiction on A/duck/Northern China/22/2017 H5 (PDB: 7DEA) of the RBS and recent mutations. Blue residues indicate those found within the 130-loop, 190-helix, or 220-loop. Red residues indicate mutations of interest. (B) Logo plots of positions 111, 199, and 214 based on geographical location. “Americas” logo plot does not include sequences from the dairy cow outbreak. (C) Amino acid alignment of HA1 from H5N1 viruses in this study. Residues in magenta are positions, 111, 199, and 214. (D) Frequency of T199 (blue) and I199 (orange) in circulating 2.3.4.4b H5N1 viruses in the Americas, including the dairy cow outbreak, between March 2022 and May 2024. (E) Maximum Clade Credibility tree of 2.3.4.4b clade H5N1 viruses in the Americas with T199 or I199 from 2022 to 2024. Posterior probabilities were marked with black dots at the nodes, with the main clades labeled by number. The size of the dots corresponds to the posterior probability values. The larger the black dot, the higher the value it represents. The scale bar at the bottom represents 0.3 substitutions per site.

### T199I is responsible for increased α2,3 sialic acid binding breadth

T199I resides in a loop on the backside of the 190-helix that leads into the 220-loop of the RBS. To determine if T199I augments binding breadth, we reverted A/Texas/37/2024 from I199 to T199 (I199T) and tested glycan binding breadth. A/Texas/37/2024 I199T demonstrated identical binding breadth to A/pelican/Chile/7087-1/2022 (**Fig. 4A-B; Extended Data Figure 1F**), indicating a mutation outside of the RBS massively affects receptor binding specificity. MD simulations predicted A/Texas/37/2024 T199 only has four different conformational states (data not shown). Furthermore, we find that T199 hydroxyl group hydrogen bonds with the amide of N248 on the same protomer, stabilizing the 190-helix and 220-loop (**Fig. 4C**). Thus, the T199I mutation would lose this stabilizing hydrogen bond, leading to more flexibility within the RBS. This additional stabilization of T199I is further emphasized by a more favorable interaction energy of T199 compared to I199 (∼-60 kcal/mol to ∼-45 kcal/mol). Analysis of the B-factor of A/Texas/37/2024 with T199 shows a decreased flexibility globally and in the 190-helix relative to A/Texas/37/2024 with the naturally occurring I199 (**Fig. 2D** and **Fig. 4D**). These data demonstrate that a single mutation outside the RBS can improve binding breadth binding breadth to distinct backbone glycans bearing terminal α2,3 sialic acids. Mechanistically, we propose that T199 stabilizes the RBS, leading to more restricted receptor binding.

**Fig. 4:**
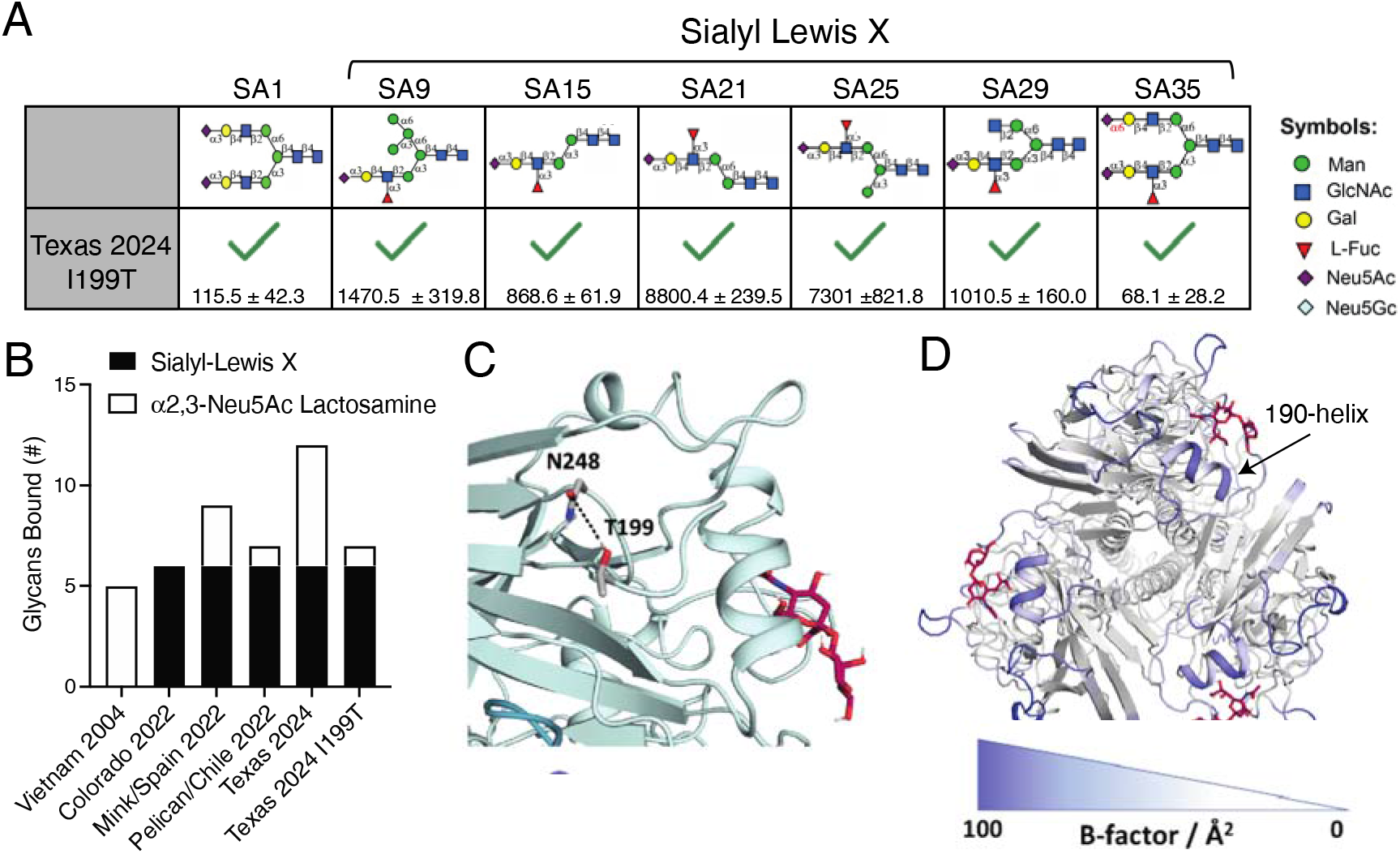
T199I is responsible for increased glycan binding breadth in A/Texas/37/2024. (A) A/Texas/37/2024 with an I199T mutation binding to distinct Neu5Ac glycans. Green checkmarks indicate a positive binding result for the corresponding glycan. Normalized relative fluorescence unit (RFU) values ± standard deviation are indicated below checkmarks. Only glycans above the background signal are shown. (B) Number and type of glycans bound by each H5. (C) Hydrogen bond analysis showing the stabilizing role of I199T with N248. (D) B-factor analysis of A/Texas/37/2024 I199T.

## DISCUSSION

Our study shows that H5N1 viruses from the ongoing dairy cow outbreak have increased their receptor binding breadth to bind more glycans bearing α2,3 sialic acids. We observed that A/Texas/37/2024 could bind 3’ sialyl Lewis X glycans, which has a fucosylated sialoside, and α2,3 sialic acid-linked lactosamine glycans. We observed a historical H5N1 virus, A/Vietnam/1204/2004, preferentially bound to α2,3 sialic acid-linked lactosamine glycans, whereas A/Colorado/18/2022, the first human case of 2.3.4.4b virus in the US, was highly specific to glycan with a 3’ sialyl Lewis X structure. A prior study found A/Vietnam/1194/2004, an isolate closely related to A/Vietnam/1204/2004, binds both 2,3 sialic acid-linked lactosamine glycans and 3’ sialyl Lewis X, albeit the former with 3-times stronger affinity^25^. Moreover, an analysis of Asian 2003-2004 H5 isolates from chickens and humans preferred sulfated α2,3 sialic acid-linked glycans, including binding to a sulfated sialyl Lewis X glycan^15^. Avian influenza viruses are known to have restricted sialic acid binding breadth as a mechanism to have specific and limited host tropism^26^. As it stands, our understanding of the glycan structures bearing α2,3 sialic acids on distinct cell types, tissues, and hosts remains poorly understood. Moreover, how host glycosylation patterns impact influenza virus evolution to augment receptor binding affinity and breadth is not well characterized. Thus, a deeper understanding of how glycan binding specificity and breadth across diverse hosts is needed to perform risk assessment of potential pandemic influenza viruses, such as H5Nx.

2.3.4.4 viruses in the mid-2010s gained mutations at positions K222Q and S227R, which increase binding to fucosylated sialosides, such as 3’ sialyl Lewis X^27^. K222 sterically clashes with the fucose group on 3’ sialyl Lewis X^27^, which could explain the selection of mutations at this site that improves binding to fucosylated glycans. Importantly, 2.3.4.4b clade H5N1 viruses have retained K222Q and S227R, which could help explain their preferential binding to 3’ sialyl Lewis X. Our data adds T199I to the list of mutations that change receptor binding, by expanding the H5 RBS binding to α2,3 sialic acid-linked lactosamine glycans. Our MD data shows that T199 stabilizes the RBS through hydrogen bonds formed with N248 on the same protomer. Since T199 is within a loop directly following the 190-helix and leads into the 220-loop, these hydrogen bonds likely stabilize the RBS, limiting the number of conformations it can adopt while binding α2,3 sialic acid-linked glycans.

Our glycan binding data shows that 2.3.4.4b H5N1 viruses, including those related to the ongoing dairy cow outbreak, have not gained binding to α2,6 sialic acids, the most abundant human receptor for influenza viruses, despite the presence of α2,6 sialic acids within the cow mammary glands^17,18^. Two mutations, E190D and G225D, are defined mutations for receptor switches between α2,3 and α2,6 sialic acid-linked glycans^22^. E190 and D190 function as direct α2,3 and α2,6 sialic acid contacts, respectively^22,28^, whereas G225D mutation introduces a bulky amino acid within the 220-loop, making binding specific to α2,6 sialic acids^29,30^. While mutations at these two sites are not observed within the circulating 2.3.4.4b, our data support that a mutation near the RBS is impacting receptor binding specificity and breadth. Notably, our data supports that mutations not directly within the RBS can dramatically change receptor binding properties. Deep mutational scanning tools for emerging influenza viruses can provide insight into mutations permissible for increased binding to α2,3 and α2,6 sialic acids^31^. A proactive analysis of H5 sequences and their potential to increase binding breadth and specificity can alert to their potential to cause a pandemic.

### Study Limitations

A limitation of our study is that we only used recombinant HA to study HA-glycan interactions, which will be limited to low avidity interactions. As a result, our approach is only detecting high affinity interactions but not low affinity high avidity interactions that would be detected using a viral particle. The glycan microarray did not include sulfated glycans, which are known to be recognized by H5Nx viruses^15^. Sulfated glycans with Neu5Ac may be an important glycan specificity of recent H5N1 viruses. Lastly, the MD data presented depend on predicted structures and serve as models of what may be occurring. Analysis of HA structures with LTSa would confirm structural confirmations taken by emerging H5N1 viruses.

## METHODS

### Sequence analyses

We downloaded 94 H5Nx sequences (Extended Data Table 2) from different clades (2.3.4.4b, 2.3.4.4c, 2.3.4.4e, 2.3.4.4g, 2.3.4.4h) along with the ancestral H5N1 sequence (A/Goose/Guangdong/1996-01-01), and two human H5N1 sequences from Asia (A/Vietnam/2004 and A/Indonesia/2005) from the GISAID database^32–34^. H5N1 avian influenza virus sequences from the 2.3.4.4b clade in the Americas were aligned using MEGA11^35,36^. The best-fit nucleotide substitution model was identified using MEGA11. A Maximum Clade Credibility (MCC) tree was constructed with BEAST v2.6.3, using the TN93+Gamma5 substitution model and partitioning by positions 1, 2, and 3^37^. The analysis employed an uncorrelated relaxed clock with a chain length of 10,000,000 generations, sampling every 1,000 generations, and discarding 10% of the samples as burn-in. The resulting file was analyzed and annotated using Tracer v1.7.1 and TreeAnnotator v1.10.4, and the annotated MCC tree was visualized using FigTree v1.4.4^38,39^. The mammal symbol on the tree originates from the Pixabay website and BioRender. Weblogo plots were generated as previously described^40^.

### Cloning and protein purification

HA sequences were downloaded from GISAID. HA ectodomains were synthesized from Integrated DNA Technologies (IDT) or Twist Biosciences and cloned into a vector with a Fibritin Foldon Domain and a his-tag. PCR-based site-directed mutagenesis was used to introduce I199T into the A/Texas/37/2024 construct. PCR reactions with mutagenesis primers were performed PrimeSTAR Max DNA Polymerase (Takara). The PCR product was treated with DpnI (New England Biolabs). All plasmids were transformed into *E. coli* New England Biolabs), miniprepped, correct clones selected, and maxiprepped. Sequence verified maxipreps were used for transfections in HEK293T cells (ATCC) or Expi293F Cells (Thermofisher). Expi293F suspension cells were maintained at 125 rpm at 37°C with 8% CO_2_ in FreeStyle^TM^293 expression medium (Gibco). HEK293T cells were grown in 37°C with 5% CO_2_. HAs were produced in-house via transfection of HEK293T cells or Expi293F cells. HAs were purified from the supernatant using nickel-NTA agarose (Qiagen) and disposable 10mL polypropylene columns (Thermofisher). HA concentrations were determined using a Pierce BCA Protein Assay (Thermofisher). HA was aliquoted and maintained at -80°C.

### Neu5Ac and Neu5Gc glycan microarray

We used a Neu5Ac and Neu5Gc glycan microarray (Zbiotech; Lot # 04242301); structures can be found here (https://www.zbiotech.com/product/neu5gc-neu5ac-n-glycan-microarray/). HA proteins were diluted into 200μL of glycan microarray assay buffer (GAAB) supplemented with 1% BSA, for a final concentration 40, 20 or 10 μg/mL of HA. Subsequently, an anti-6x His tag antibody and anti-rabbit immunoglobulin (H+L) (Cy3) antibody were added to the HA + GAAB mixture at a final concentration of 3.2 μg/mL each. The entire mixture was then incubated at room temperature for 60 minutes with gentle vortexing as part of the precomplexing process. To prepare the microarray slide for analysis, the slide was pretreated with glycan microarray blocking buffer (GABB) supplemented with 1% BSA at room temperature for 60 minutes. Following this, the precomplexed HA protein samples were added to the microarray, with 100μL added to each submicroarray. The slide was incubated for 60 minutes at room temperature to facilitate binding interactions. After this incubation period, the slide was thoroughly washed to remove any unbound components. The slide was then scanned at 532 nm using high intensity (1 PMT) to detect and visualize any interactions. Innopsys’ Mapix software was used to analyze the microarray scans. Positive binding signals were determined by subtracting the background and negative control signals from all experimental sample signals.

### MD simulations

The starting structures of A/Colorado/18/2022, A/Texas/37/2024, and A/Texas/37/2024 I199T, were predicted using *ColabFold*, which increased the accessibility of protein structure prediction tools by combining *AF2* with the rapid homology search capability of MMseqs2, making it an easy to use and fast software (∼90-fold speed up in prediction) to predict homo-and heteromeric complexes, matching the prediction quality of AF2 and AF-multimer^41,42^. As template to model the sialic acid complex (LSTa) we used the available X-ray structure of A/California/04/2009 in complex with LSTa (PDB ID: 3UBJ).

In addition, we used the GlycoShape tool to ensure the known glycosylation sites are glycosylated for the simulations^43^. For our simulations we capped the C-terminal and N-terminal parts of each domain with acetylamide and N-methylamide to avoid perturbations by free charged functional groups. For each H5 variant, we performed three repetitions of 500 ns of classical molecular dynamics simulations using the AMBER 22 simulation software package which contains the pmemd.cuda module^44^. The structures were prepared using CHARMM-GUI^45,46^. The structure models were placed into cubic water boxes of TIP3P water molecules^47^ with a minimum wall distance to the protein of 12 Å^48,49^. Parameters for all simulations were derived from the AMBER force field 14SB^50,51^. To neutralize the charges, we used uniform background charges^52–54^. Each system was carefully equilibrated using a multistep equilibration protocol^55^. Bonds involving hydrogen atoms were restrained using the SHAKE algorithm, allowing a timestep of 2.0 femtoseconds^56^. The systems’ pressure was maintained at 1 bar by applying weak coupling to an external bath using the Berendsen algorithm^57^. The Langevin Thermostat was utilized to keep the temperature at 300K during the simulations^58^.

### MD analysis

For all investigated H5 variants, we calculated the respective contacts of the HA protomers with LSTa in solution using the GetContacts software (Stanford University; https://getcontacts.github.io/). This tool can compute interactions within one protein structure, but also between different protein interfaces and allows to monitor the evolution of contacts during the simulation. Apart from visualizing and quantifying the contacts of the different poses, we calculated the residue-wise B-factor, as measure of global flexibility implemented in cpptraj^59^ to identify differences in the conformational diversity between the HA variants. We used PyMOL to visualize protein structures (PyMOL - The PyMOL Molecular Graphics System, Version 3.0 Schrödinger, LLC).

### HA modeling

The protein structure 7DEA, from A/duck Northern China/22/2017 (H5N6), was retrieved from the Protein Data Bank (PDB) and visualized using PyMOL (Version 2.6, Schrödinger, LLC). All numbering in this manuscript is H3-numbering, based on Burke and Smith^60^.

## ACKNOWLEDGEMENTS

We gratefully acknowledge all data contributors, including the authors and their originating laboratories responsible for obtaining the specimens, and their submitting laboratories for generating the genetic sequence and metadata and sharing via the GISAID Initiative, on which this research is based. We also thank Jian Zheng, Xi Chen, and Nick Berning of Z Biotech for their discussion and suggestions on glycan analyses. We thank Julianna Han for her critical feedback on data interpretation.

## AUTHOR CONTRIBUTIONS

Conceptualization: MRG, WJ, JJG; Methodology: MRG, WJ, MLF-Q, JJG; Investigation: MRG, WJ, MLF-Q, JJG; Visualization: MRG, WJ, MLF-Q, JJG; Funding acquisition: JJG and ABW; Project administration: JJG; Supervision: JJG and ABW; Writing – original draft: MRG, WJ, JJG; Writing – review & editing: MRG, WJ, MLF-Q, ABW.

---

**Extended Data Figure 1:**
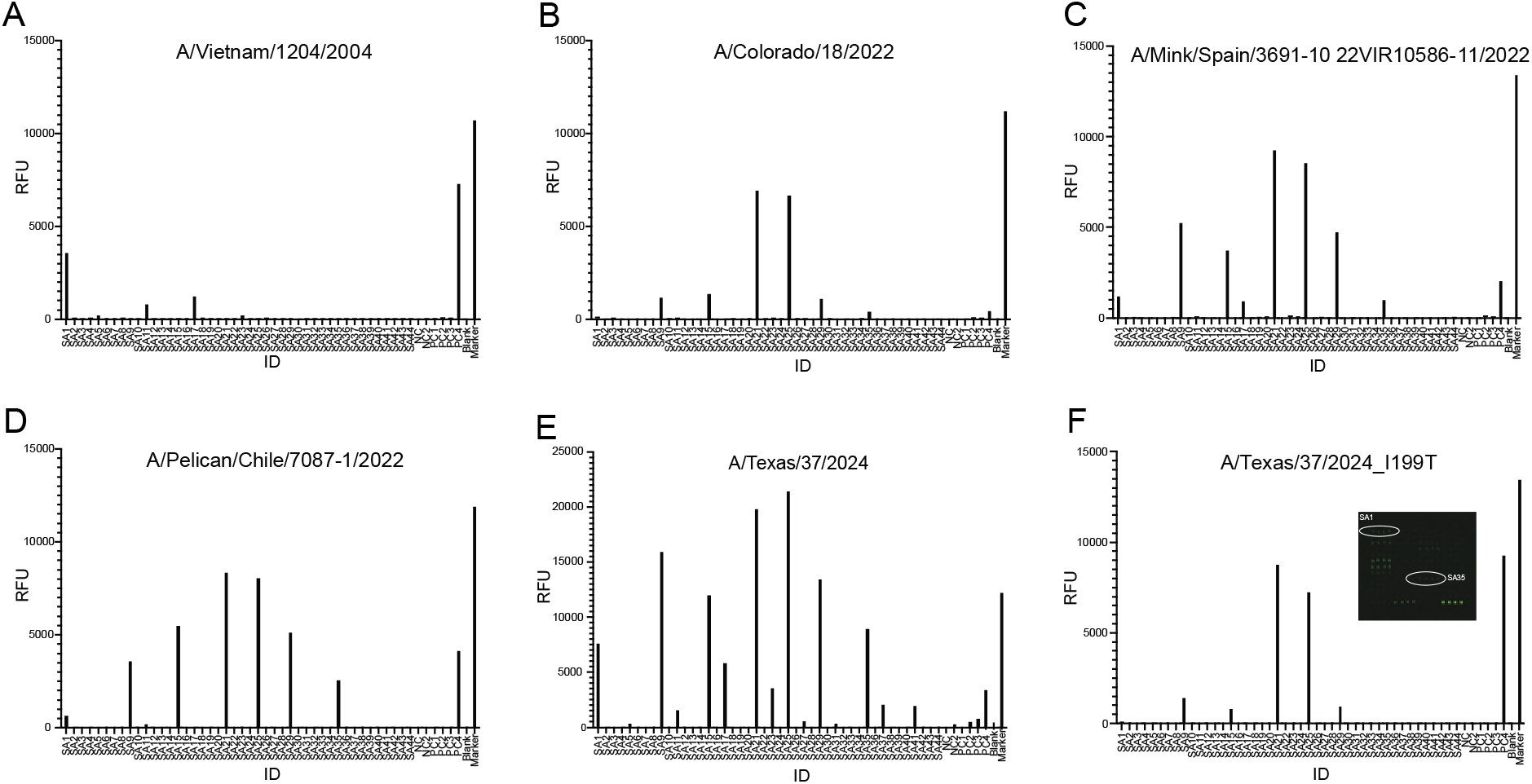
rHA binding to glycan microarray. (A-F) rH5 binding RFUs for each glycan on the microarray. rH5s tested are A/Vietnam/1204/2024 (A), A/Colorado/18/2022 (B), A/Mink/Spain/3691-1022VIR10586-11/2022 (C), A/Pelican/Chile/7087-1/2022 (D), A/Texas/37/2024 (E), and A/Texas/37/2024 I199T (F). Microarray fluorescence is shown in F, with SA1 and SA35 circled to indicate detection of fluorescence despite low RFU values.

**Extended Data Table 1**: The HA sequences used for the phylogenetic tree analysis.

**Extended Data Table S2:** Glycans in the Neu5Ac and Neu5Gc microarray.

